# CRISPR-Cas-Docker: Web-based *in silico* docking and machine learning-based classification of crRNAs with Cas proteins

**DOI:** 10.1101/2023.01.04.522819

**Authors:** Ho-min Park, Jongbum Won, Yunseol Park, Esla Timothy Anzaku, Joris Vankerschaver, Arnout Van Messem, Wesley De Neve, Hyunjin Shim

**Author notes:** Corresponding Author: Hyunjin Shim.

## Abstract

**Motivation:** CRISPR-Cas-Docker is a web server for *in silico* docking experiments with CRISPR RNAs (crRNAs) and Cas proteins. This web server aims at providing experimentalists with the optimal crRNA-Cas pair predicted computationally when prokaryotic genomes have multiple CRISPR arrays and Cas systems, as frequently observed in metagenomic data. CRISPR-Cas-Docker provides two methods to predict the optimal Cas protein given a particular crRN sequence: a structure-based method (*in silico* docking) and a sequence-based method (machine learning classification). For the structure-based method, users can either provide experimentally determined 3D structures of these macromolecules or use an integrated pipeline to generate 3D-predicted structures for *in silico* docking experiments.

**Results:** CRISPR-Cas-Docker is an optimized and integrated platform that provides users with 1) 3D-predicted crRNA structures and AlphaFold-predicted Cas protein structures, 2) the top-10 docking models for a particular crRNA-Cas protein pair, and 3) machine learning-based classification of crRNA into its Cas system type.

**Availability and implementation:** CRISPR-Cas-Docker is available as an open-source tool under the GNU General Public License v3.0 on GitHub. It is also available as a web server.

## 1. Introduction

CRISPR-Cas is a prokaryotic adaptive immune system [1,2] that consists of two genetic components: (1) CRISPR arrays with CRISPR RNAs (crRNAs) encompassing short palindromic repeats and unique spacers from previous infections and (2) CRISPR-associated systems (Cas) which form a complex of proteins to cleave invading foreign genetic elements. CRISPR-Cas systems have been repurposed as genome-editing tools [3,4] and antimicrobials [5,6], with this biotechnological potential driving the scientific community to discover novel types of CRISPR-Cas systems [7–9].

CRISPR arrays are assumed to be associated with Cas systems when they are co-located in prokaryotic genomes (usually within ±10,000 base pairs). However, metagenomic data from diverse environments have revealed that prokaryotic genomes often have multiple CRISPR arrays and Cas systems. Such complexity in genomic architecture can lead to suboptimal RNA-protein interactions between the crRNA-Cas protein complex in CRISPR-Cas-based genomic tools [10]. In a previous study, we predicted crRNAs that bind optimally to a particular Cas protein through *in silico* docking experiments, suggesting that such *in silico* experiments can be adopted as a preliminary approach to design stable CRISPR-based antimicrobials using the newly discovered Cas13 proteins [11].

Here, we present a web application named CRISPR-Cas-Docker that offers an optimized and integrated pipeline to conduct *in silico* docking experiments between a crRNA and a Cas protein (Fig. S1). By leveraging our expertise with RNA structure prediction, AlphaFold-based protein structure prediction, and *in silico* macromolecular docking, we aim at providing experimentalists with a practical and user-friendly bioinformatics tool that can suggest the most optimal crRNA-Cas protein pairs to be tested *in vitro*.

## 2. Implementation

### 2.1 Predicting the 3D structures of crRNAs and Cas proteins

*In silico* docking requires the availability of the 3D structures of biological macromolecules, which can be obtained through experimental techniques such as X-ray crystallography, NMR, and cryoelectron microscopy [12]. If experimentally determined structures are not available, these 3D structures can be estimated rapidly and accurately through (1) deep learning-based protein structure prediction programs such as AlphaFold [13,14] and (2) a combination of 2D and 3D RNA structure prediction programs [15,16]. Using the experimentally determined structures of Cas proteins, we verified that AlphaFold is able to achieve an adequate level of prediction accuracy for large effector proteins such as Cas13 (Table S1). We used AlphaFold to fold four Cas13 proteins with and without a template. The average (standard deviation) of the TM-score was 0.992 (0.001) and 0.817 (0.012), with and without a template, respectively. CRISPR-Cas-Docker has an integrated option to generate a 3D-predicted RNA structure and an AlphaFold-predicted protein structure for a crRNA sequence and a Cas protein sequence, respectively (Fig. 1a, b).

**Fig. 1:**
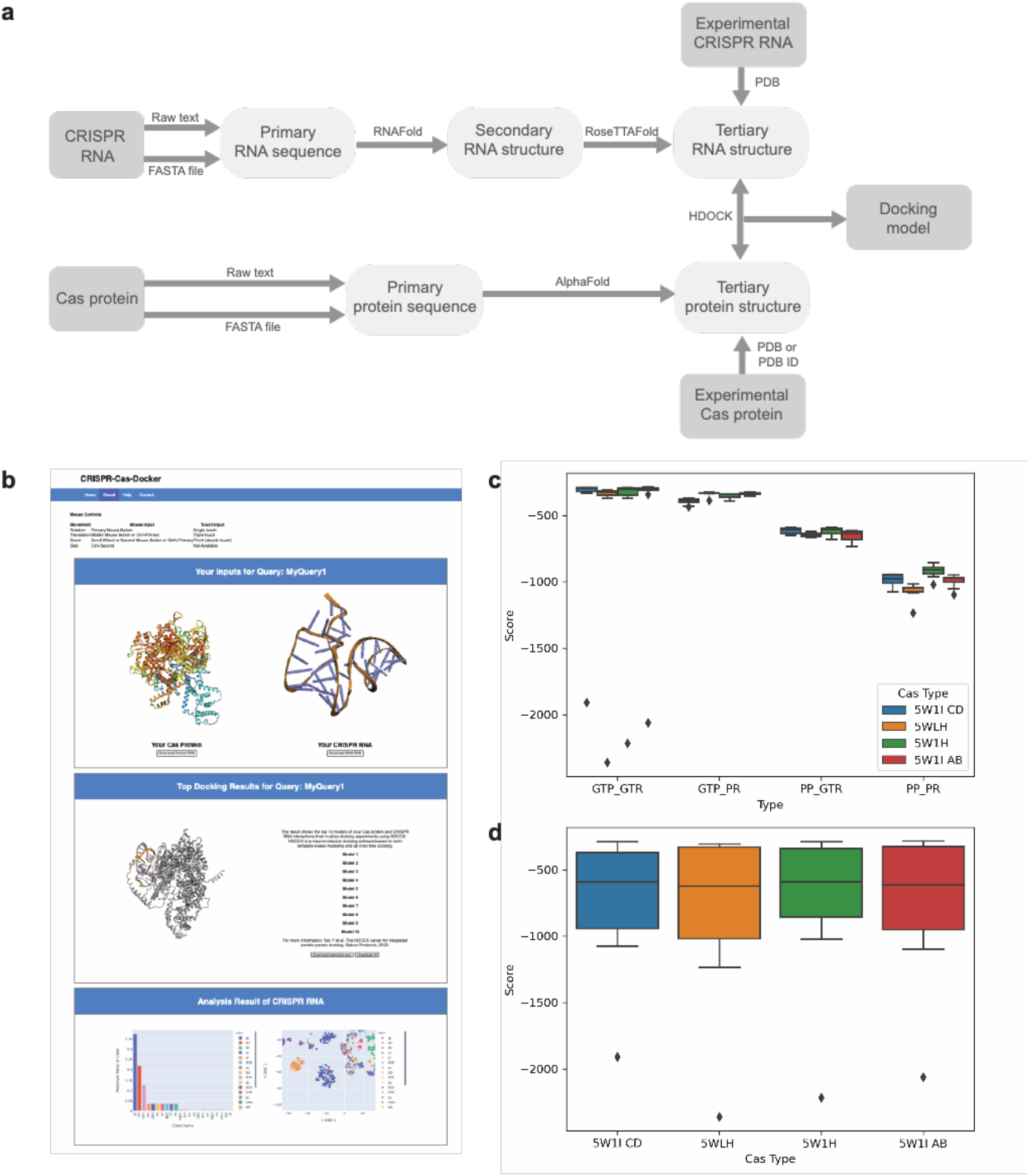
CRISPR-Cas-Docker. **(a)** Workflow used by CRISPR-Cas-Docker. **(b)** Results page generated by CRISPR-Cas-Docker, showing the downloadable PDB files of an AlphaFold-predicted Cas protein structure, a 3D-predicted crRNA structure, and the top-10 docking models. **(c)** Performance of CRISPR-Cas-Docker, using individual boxplots to show the docking scores obtained for different Cas13 proteins. **(d)** Performance of CRISPR-Cas-Docker, showing the distribution of docking scores obtained for different types of Cas proteins with GTP and PP combined. According to the HDOCK server, a lower docking score indicates a better docking model. (GTP: Ground Truth Cas Protein; GTR: Ground Truth crRNA; PP: Predicted Cas Protein (AlphaFold); PR: Predicted crRNA (RoseTTAFold)).

### 2.2 *In silico* docking of crRNAs and Cas proteins

In earlier work, we determined the best program to conduct *in silico* experiments between crRNAs and Cas proteins to be HDOCK [17], leading to the best RNA-protein docking and binding affinity results using an experimentally validated dataset [11]. CRISPR-Cas-Docker uses the template-free docking approach of HDOCK to generate the top-10 docking models for a given crRNA-Cas protein pair, with the docking score of each model calculated by statistical mechanics-based energy scoring functions [18]. Previously, we verified that a docking score is a strong indicator of the binding affinity between crRNA-Cas protein complexes [11]. We compared the docking scores between all combinations of experimentally determined and computationally predicted crRNAs and Cas proteins (Fig. S2). According to this performance study, AlphaFold-predicted proteins docked equally well or even better with the experimental crRNA and the 3D-predicted crRNA (Fig. 1c, d). From these results, we conclude that the effectiveness of docking is not affected by the use of predicted structures instead of experimental structures. The final step of CRISPR-Cas-Docker requires human expertise to identify the best *in silico* docking model from the generated top-10 docking models, using biological information such as the location of binding sites and the orientation of bound crRNA.

### 2.3 Machine learning-based classification of crRNAs

CRISPR-Cas-Docker includes support for machine learning-based classification of an input crRNA sequence into its associated Cas system type [7–9]. This feature is a sequence-based prediction of the optimal Cas protein for a particular crRNA sequence, which is an alternative method to the structurebased prediction of optimal crRNA-Cas pairs. To learn the associations between CRISPR arrays and Cas systems, we first created a dataset of CRISPR arrays labeled with their co-localized Cas system type (Fig. S3-S7). To that end, we extracted the CRISPR-Cas systems from the CRISPRCasdb [19] and labeled the CRISPR arrays co-localized within ± 10,000 base pairs with their corresponding Cas system (Table S2). Next, on the curated dataset, we trained a supervised machine learning method that is known as K-Nearest Neighbors (KNN) [20]. Although KNN is one of the simplest classifiers in the area of machine learning, it has been used widely in the fields of gene and protein prediction, thanks to its interpretability, even when making use of complex data [21–24]. The classification analysis shows an overall prediction accuracy of 92.3%, confirming the ability of KNN to act as an accurate and efficient classifier of crRNAs into their associated Cas system type (Table S3, Figure S8).

## 4. Conclusion

Designed for experimental biologists, CRISPR-Cas-Docker addresses the need to predict optimal crRNA-Cas protein pairs *in silico* before conducting expensive and time-consuming experiments. As metagenomic data become widely available, this bioinformatics tool enables performing a rapid preliminary study to disentangle the complex associations between multiple CRISPR arrays and Cas systems in prokaryotic genomes. Currently, CRISPR-Cas-Docker produces 3D-predicted structures of crRNAs and Cas proteins, top-10 docking models, and interactive graphs to visualize the machine learning-based classification of an input crRNA into its Cas system type. As future prospects, we aim at integrating AlphaFold-Multimer as a protein prediction program, making it possible to have Cas proteins with multi-unit effectors as an input to CRISPR-Cas-Docker.

## Data and Code Availability

2D RNA structures and 3D RNA structures were predicted with ViennaRNA v.2.5.1 and RoseTTAFold v.2.0.0, respectively. *In silico* docking experiments were performed with HDOCK v.1.1.0. Protein structures were predicted with AlphaFold2, available under an open-source license at https://github.com/deepmind/alphafold. As protein structure similarity metrics, we used TM-align (https://zhanggroup.org/TM-align). 3-D structure visualizations were created with 3Dmol.js (https://3dmol.csb.pitt.edu/doc/tutorial-embeddable.html). For data analysis purposes, Python 3.8.13 (https://www.python.org), NumPy v.1.23.4 (https://github.com/numpy/numpy), Seaborn v.0.12.0 (https://github.com/mwaskom/seaborn), Matplotlib v.3.5.3 (https://github.com/matplotlib/matplotlib), and Pandas v.1.4.3 (https://github.com/pandas-dev/pandas) were used.

## Acknowledgments

We would like to thank Jill Banfield for inspiring the machine learning-based classification of CRISPR RNAs into their Cas system types. Furthermore, the research and development activities described in this study were funded by Ghent University Global Campus (GUGC), Incheon, Korea.

## Supplementary Information

Supplementary data will be provided upon request.

