## Supplemental figures and tables for "CRISPR-Cas-Docker: Web-based *in silico* docking and machine learning-based classification of crRNAs with Cas proteins"

### Author Information

#### Affiliations

[1]**Center for Biosystems and Biotech Data Science, Ghent University Global Campus, Incheon 21985, South Korea**

[2]**Department of Electronics and Information Systems, Ghent University, Ghent 9000, Belgium**

[3]**Department of Applied Mathematics, Computer Science and Statistics, Ghent University, Ghent 9000, Belgium**

[4]**Department of Mathematics, University of Liège, Liège 4000, Belgium**

**Figure S1. The architecture of CRISPR-Cas-Docker.** This diagram shows how users interact with the Server to make service requests and view results. The *Server* manages these interactions through the *Home* and *Result* interfaces. The *Worker* component is responsible for generating the actual results, with the *Server* and *Worker* exchanging data through Storage without direct communication. The black arrow represents the request of a user, while the red arrow shows the generation of results. The blue arrow indicates the user interaction with the results. CRISPR-Cas-Docker is implemented by making use of the following Python libraries or binaries;

Server: Flask, BioPython, Plotly, NumPy, Pandas

Workers: Scikit-learn, RNAfold, RoseTTAFold, AlphaFold, HDOCK.

*The source of the libraries and binaries can be found in the section on Data and Code Availability of the main manuscript.

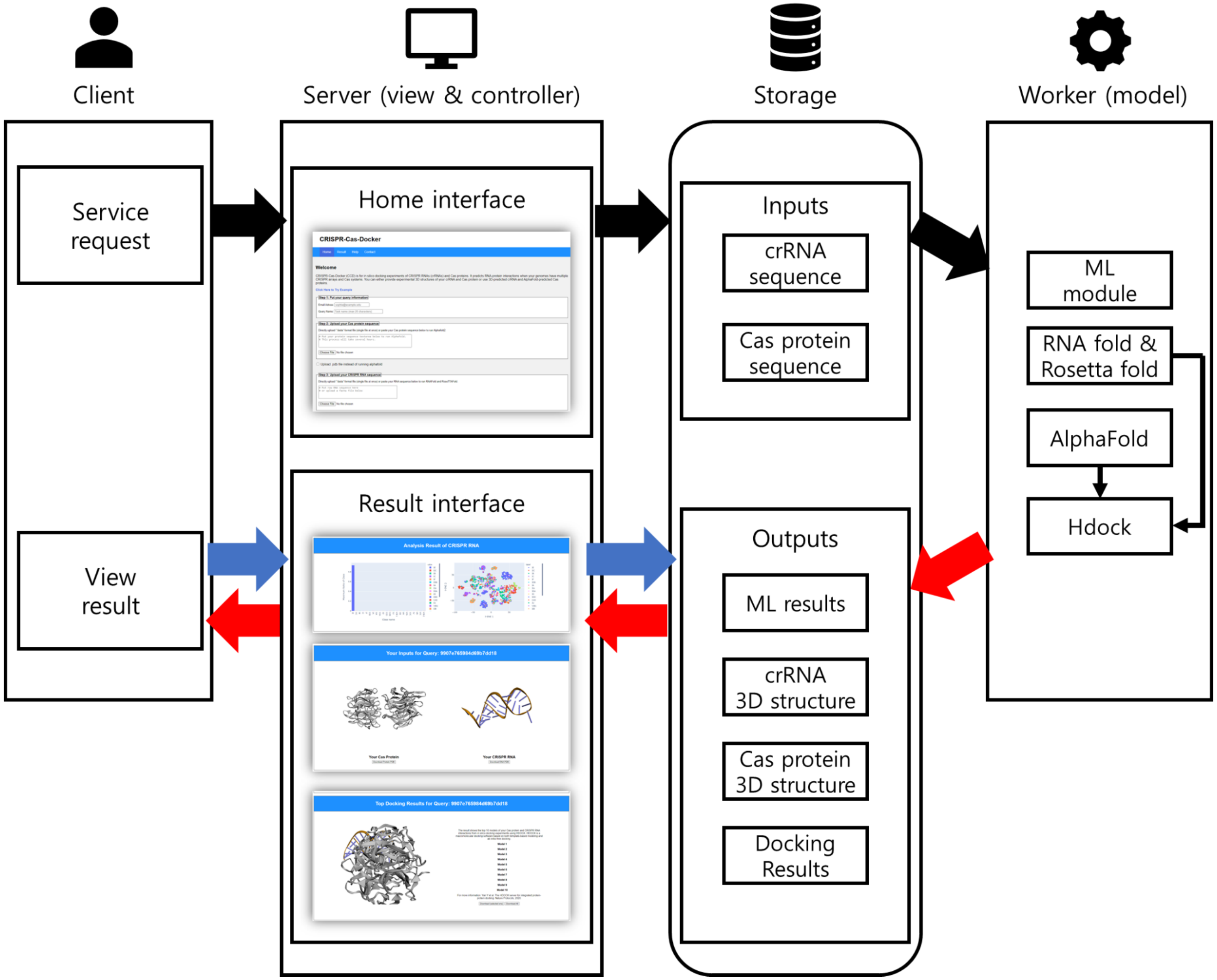

**Figure S2.** Averaged boxplot of CRISPR-Cas-Docker performance for Cas13 proteins. In particular, this boxplot shows that the average docking score is approximately -600 for all four Cas proteins, with no noticeable differences between them. However, there are some particularly low outliers for GTP-GTR, which may be indicative of docking performance very close to the ground truth. According to the HDOCK server, a lower docking score corresponds to a better docking model. (GTP: Ground Truth Cas Protein; GTR: Ground Truth crRNA; PP: Predicted Cas Protein (AlphaFold); PR: Predicted crRNA (RoseTTAFold)).

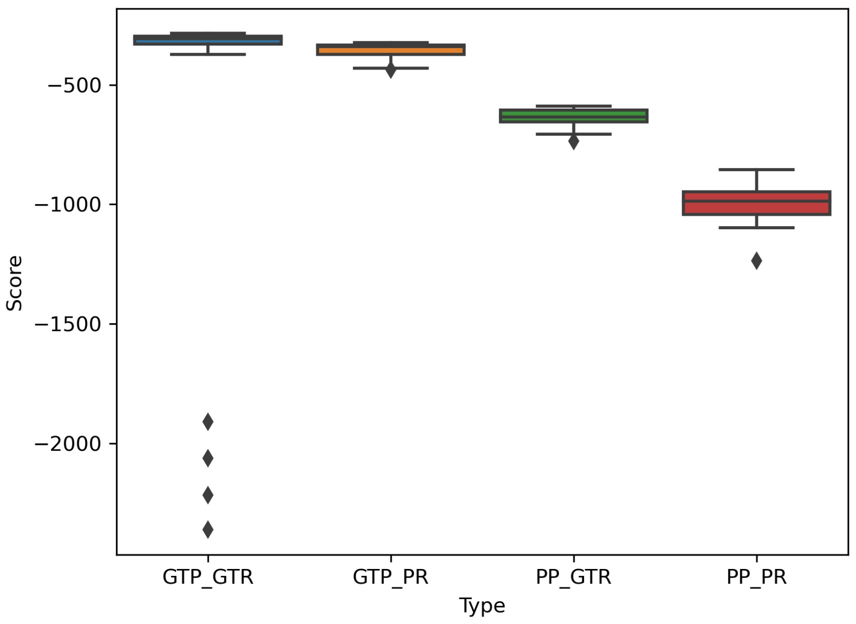

**Figure S3. CRISPR repeat sequences labeled by the adjacent Cas system** (±10,000 base pairs). As the created dataset has a class imbalance, we divided the CRISPR repeat sequences into four subsets based on their frequency of occurrence. This ensures that the KNN classification is not affected by the aforementioned class imbalance. The first subset, which is named Major, includes IE, IIC, IB, IC, IF, and IIIA, with each class containing more than 1,000 instances. Since the number of IE instances (6,862) is four or more times that of other types, 20% of the IE repeats were randomly sampled for training (1,372). The second subset, which is called Minor, includes IIIB, IA, IIA, and IIID, with each class having more than 300 instances and not belonging to Major. The third subset is named Tiny, which includes classes with less than 300 instances and with these classes not belonging to either Major or Minor. Lastly, the subset Undefined consists of CAS and IU which are Cas system types that are not complete and unidentified, respectively.

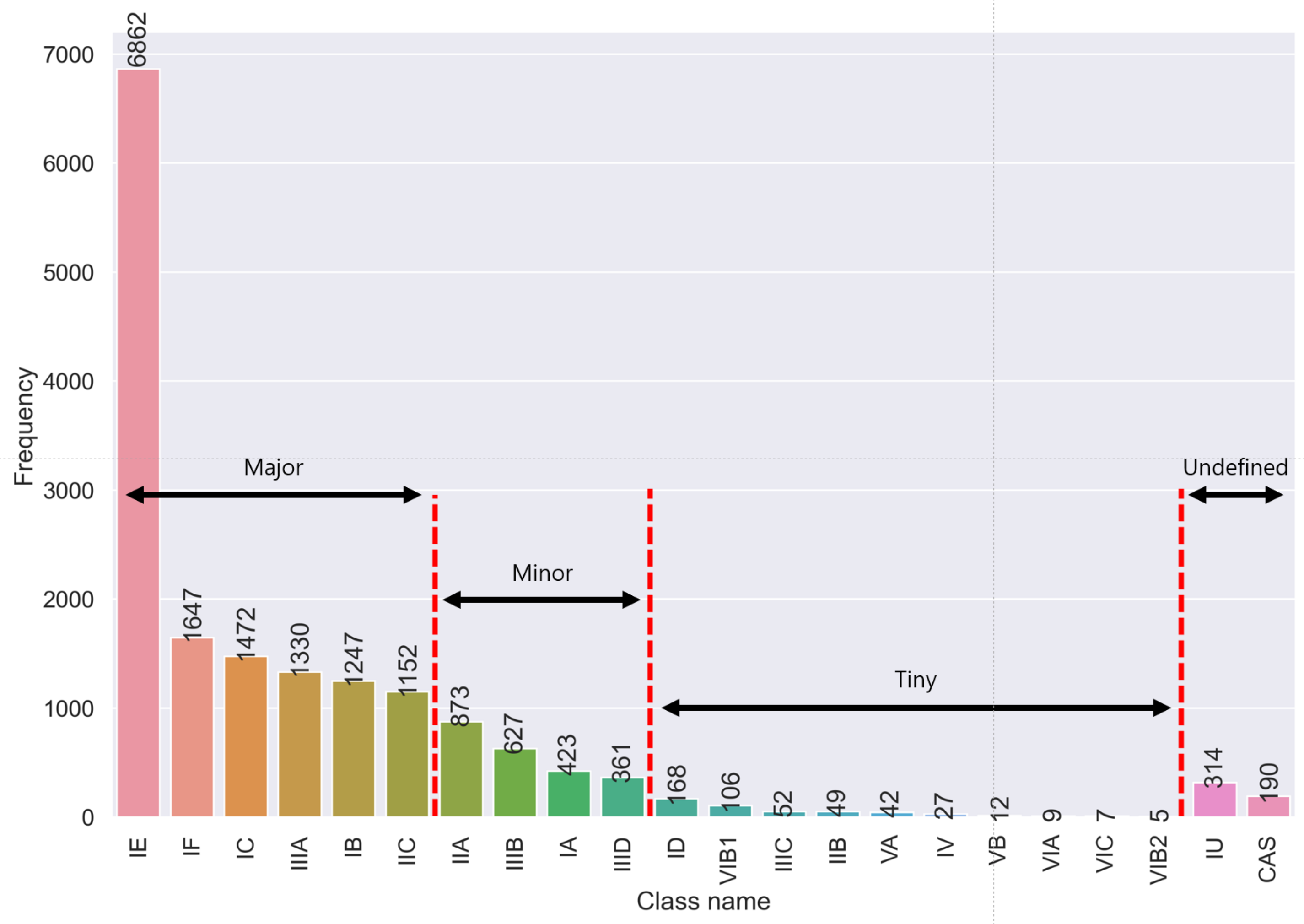

**Figure S4. Distance distributions of CRISPR repeat sequences labeled with their adjacent Cas system.** This histogram shows the distance of each CRISPR array to the adjacent Cas system in base pairs. It shows that most CRISPR arrays are around 100 base pairs away from their adjacent Cas system (2859), but there are some as far as 10,000 base pairs away.

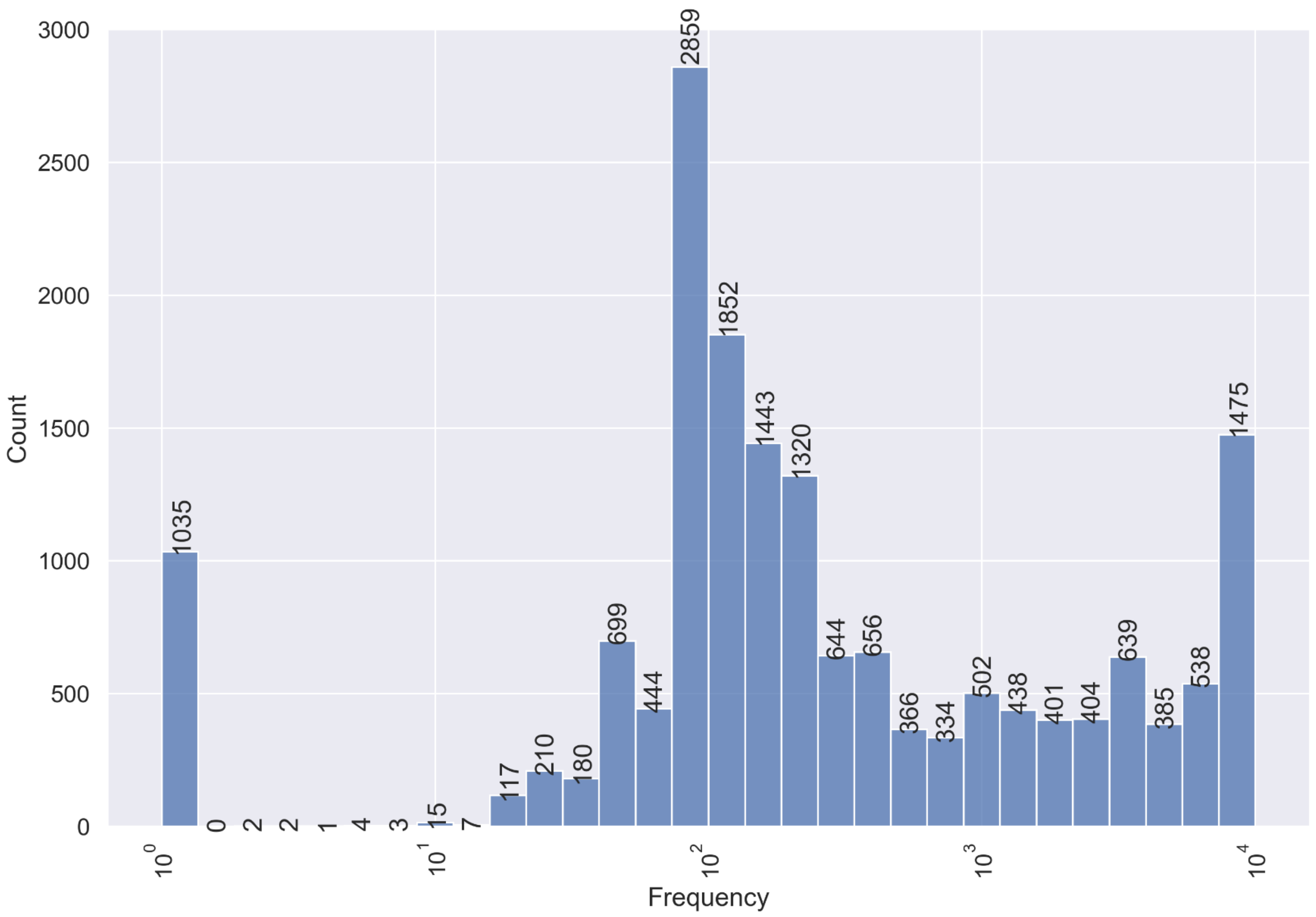

**Figure S5. Length distribution of CRISPR repeat sequences labeled with their adjacent Cas system** (±10,000 base pairs). The number next to the type of CRISPR repeats at the top of each histogram shows the average length (standard deviations), which indicates that the average length of CRISPR repeats varies by the associated Cas system.

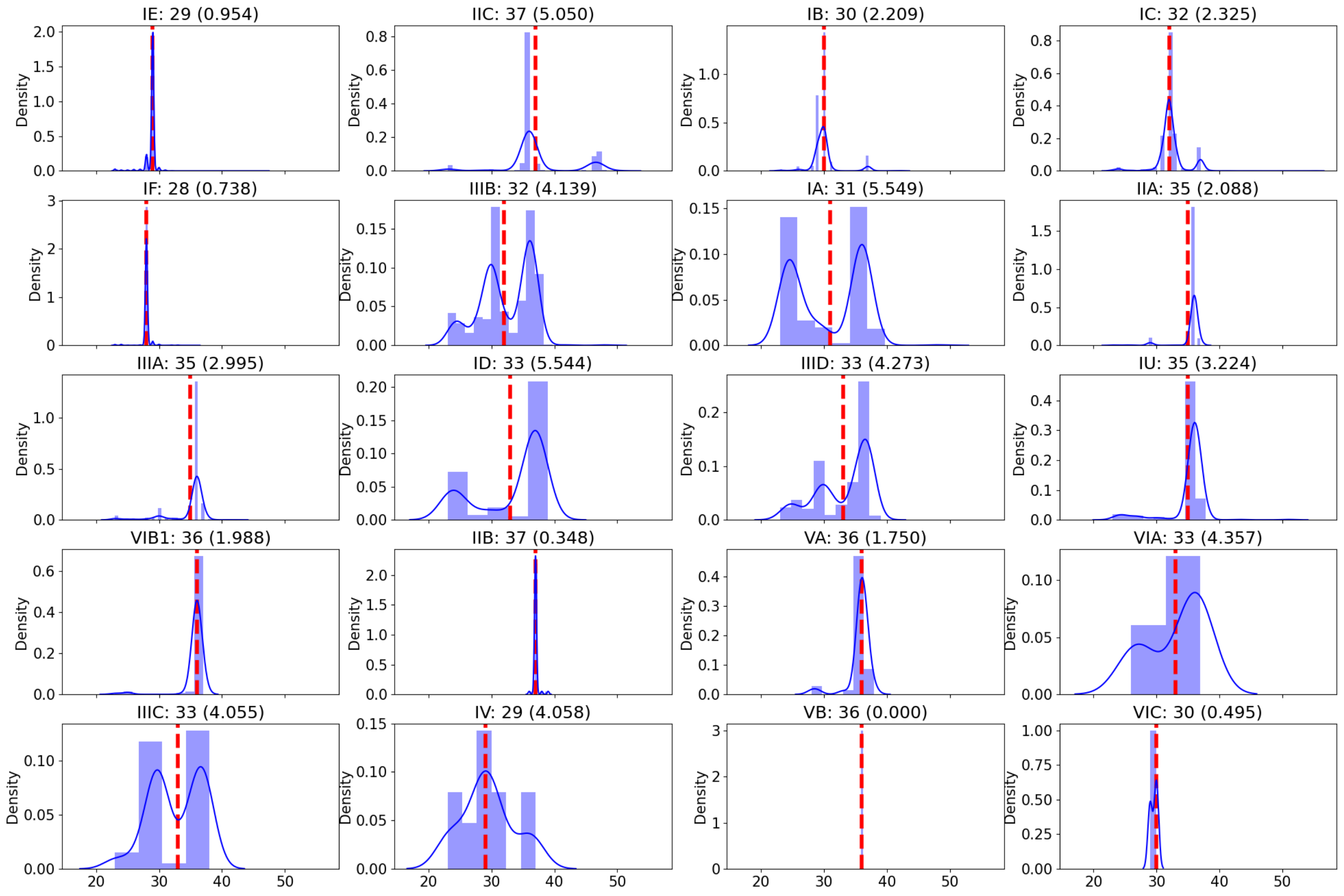

**Figure S6. Two-dimensional CRISPR sequence atlas.** The interactive version is available in CRISPR-Cas-Docker. We used t-SNE to show the Hamming distance between all pairs of sequences in a two-dimensional representation. Each dot represents a crRNA sequence, with the shape and color of the dot indicating the type of that particular crRNA sequence. According to the t-SNE method, closely located dots denote similar sequences. We pad the shorter sequence with padding characters in order to equalize their lengths when using the Hamming distance measure.

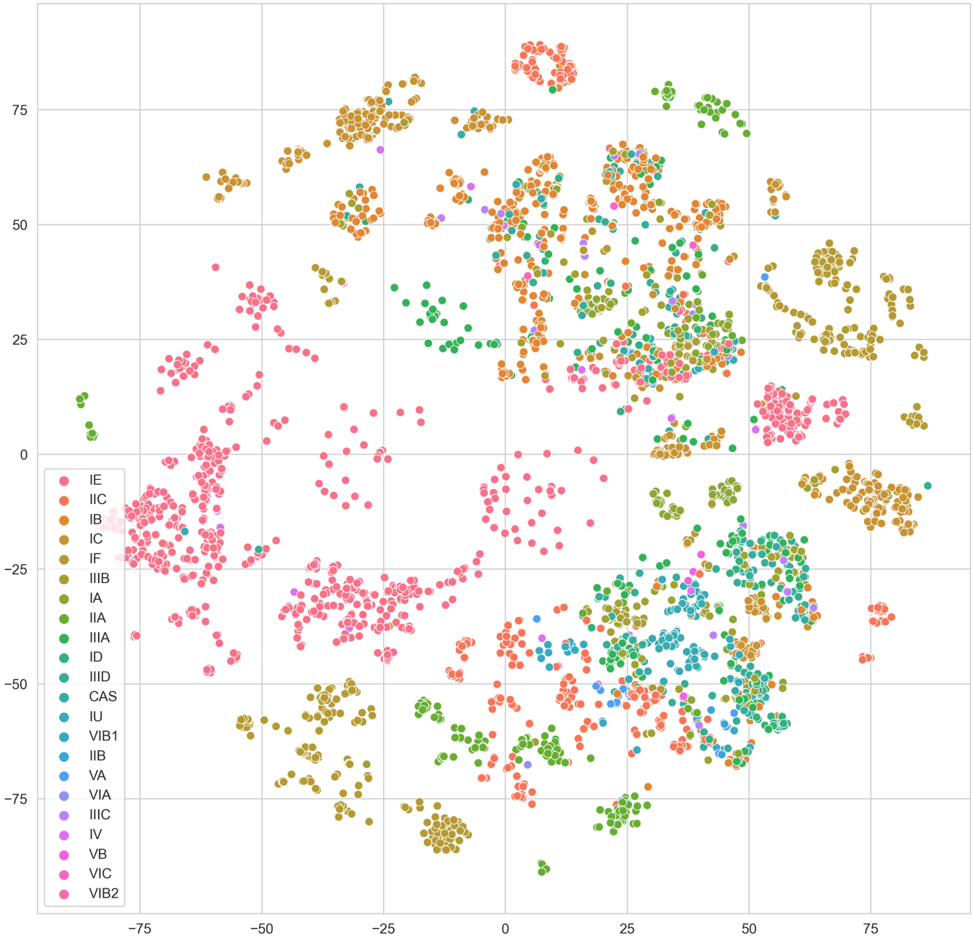

**Figure S7. Two-dimensional CRISPR sequence atlas, separated by the data subsets.** We divide the crRNA sequence data into (a) Major (more than 1,000 sequences), (b) Minor (more than 300 sequences), (c) Tiny (less than 300 sequences), and (d) Undefined (CAS and IU types). In the case of the Major subset, we found that the IE type has four clusters, and the cluster located near (0, 25) overlaps heavily with other types. In the case of the Minor subset, we found many overlapping points in most of the types, except for IIID. These overlapping points suggest that a single crRNA sequence may be labeled with multiple Cas system types.

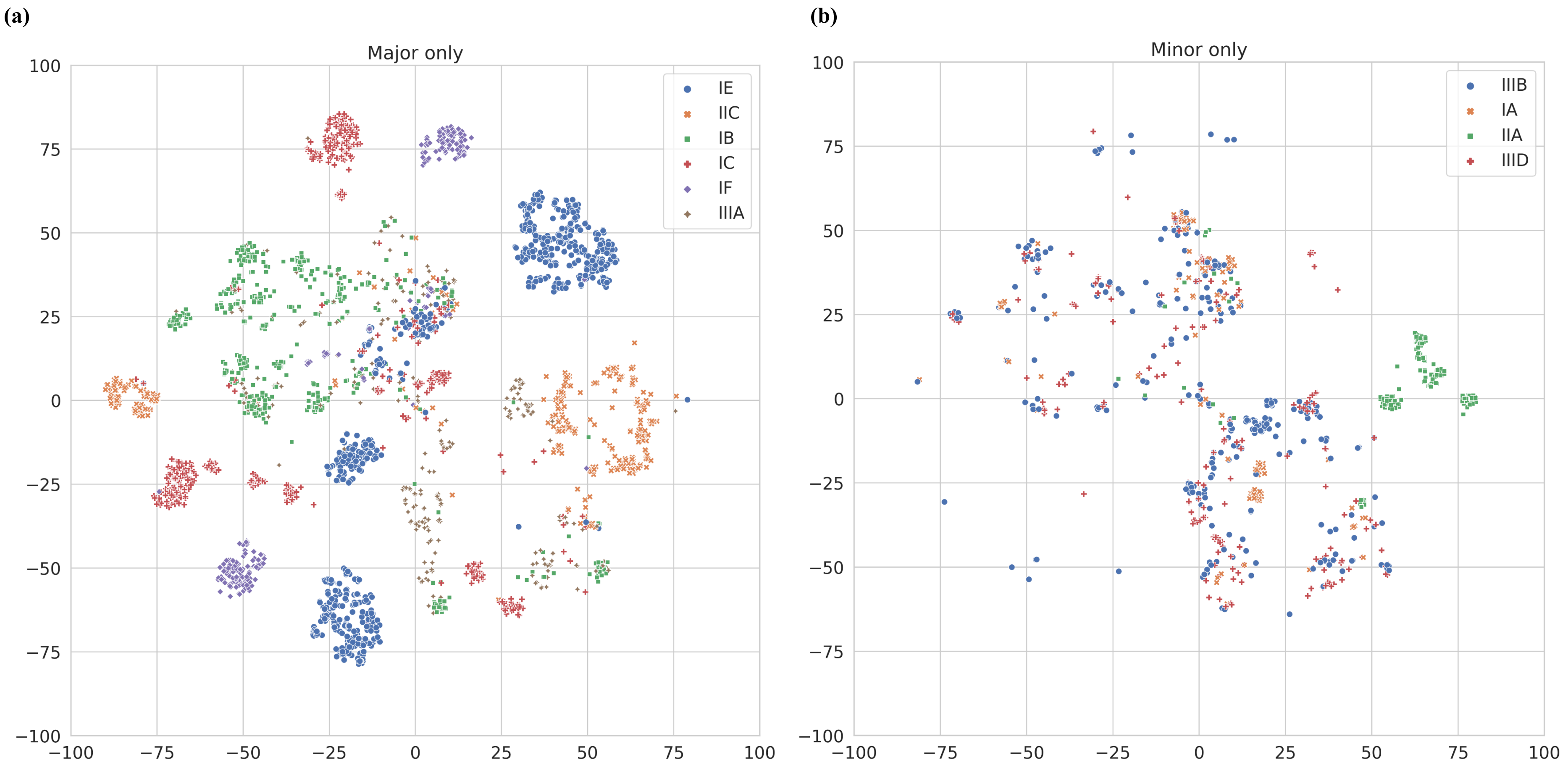

(c)

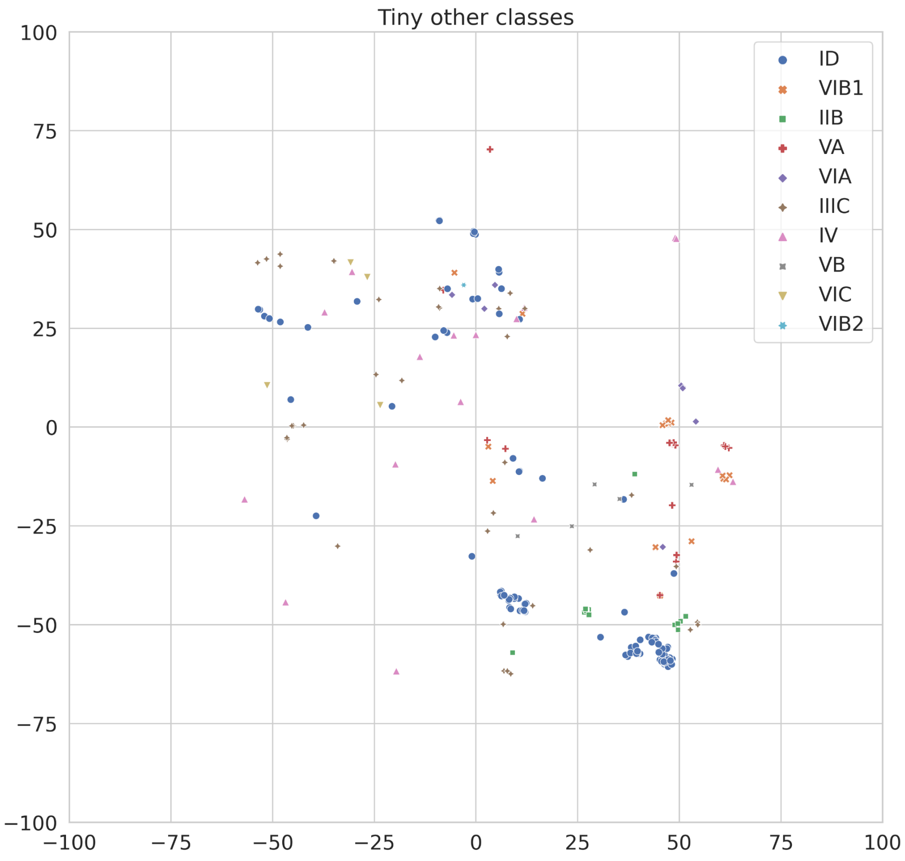
Tiny other classes
ID
VIB1
IIB
VA
VIA
IIIC
IV
VB
VIC
VIB2
100
75
50
25
0
−25
−50
−75
−100
−100
−75
−50
−25
0
25
50
75
100

(d)

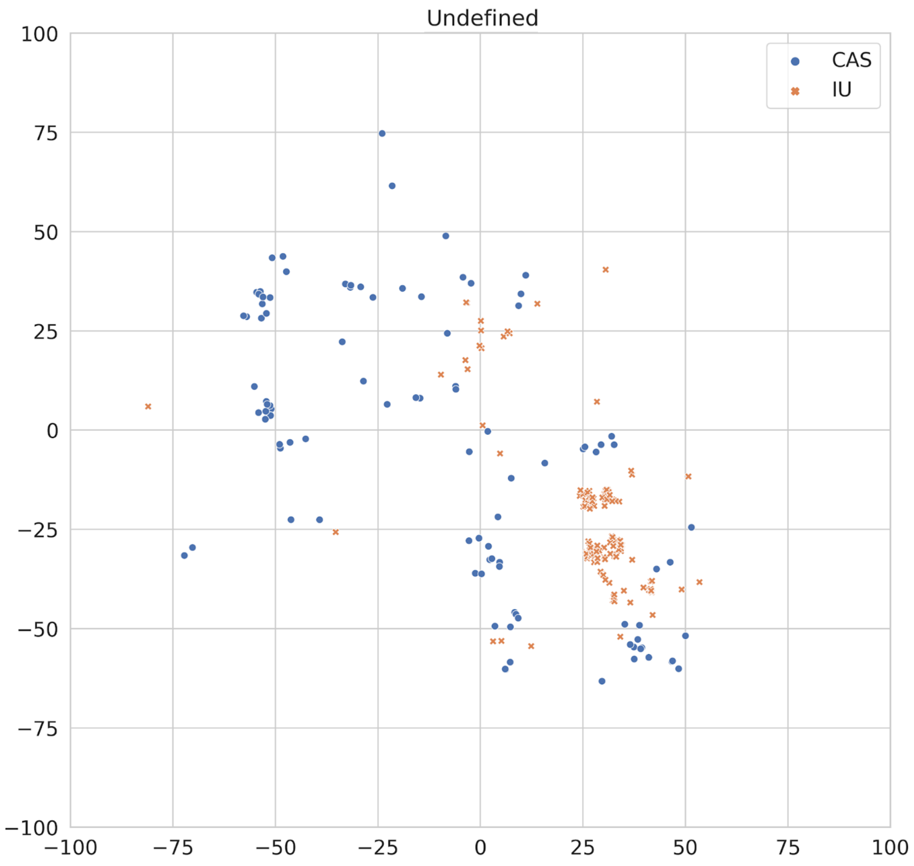
Undefined
CAS
IU
100
75
50
25
0
−25
−50
−75
−100
−100
−75
−50
−25
0
25
50
75
100

**Figure S8. Confusion matrix of the classification result.** The matrix is divided into four subsets for easier viewing. Except for the Tiny subset, which has a small number of test sets, IB had the highest misclassification rate in the Major subset. We observed that IB is frequently mistaken for IIIB. In the Minor subset, IIIB had the highest misclassification rate. This may be due to IIIB sharing many similar sequences with other CRISPR arrays. Overall, the KNN model performed well in correctly classifying the Major and Minor classes, with the aforementioned exceptions.

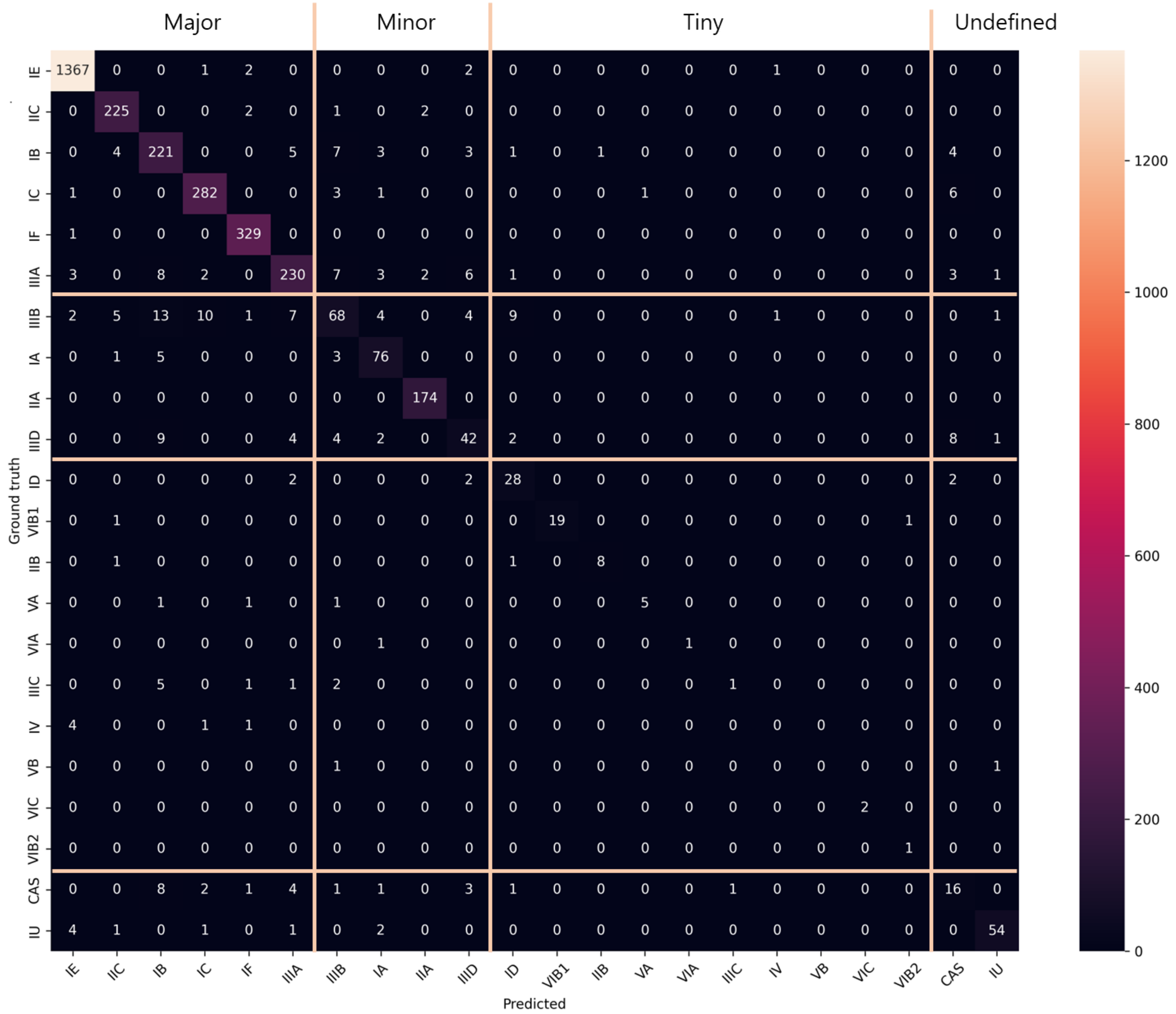

**Table S1: TM-scores of AlphaFold-predicted Cas proteins.**

TM-score (0,1] to measure the folding performance of AlphaFold.
 0.0 < TM-score < 0.30, random structural similarity
 0.5 < TM-score ≤ 1.00, high structural similarity (about the same fold)

RMSD to check the atom level structure difference.
RMSD < 2Å: two structures are the same structure.

| TM-Score | 5w1h | 5w1i_AB | 5w1i_CD | 5wlh |
|---|---|---|---|---|
| **With template** | 0.99390 | 0.99240 | 0.99077 | 0.99403 |
| **No template** | 0.80226 | 0.83607 | 0.81096 | 0.81735 |

| RMSD | 5w1h | 5w1i_AB | 5w1i_CD | 5wlh |
|---|---|---|---|---|
| **With template** | 1.18 | 1.23 | 1.31 | 1.31 |
| **No template** | 4.07 | 4.03 | 4.00 | 4.40 |

**Table S2. Distribution of CRISPR array evidence level by type.** The numbered suffix (1 to 4) indicates the evidence level, which is assigned based on the combined degree of similarity of repeats and spacers (Couvin et al. 2018). A higher evidence level indicates a higher chance that the sequence corresponds to a CRISPR array.

| Cas with evidence level | Frequency | Percentage | | Cas with evidence level | Frequency | Percentage |
|---|---|---|---|---|---|---|
| Cas-Type IE 1 | 143 | 0.021 | | CAS 1 | 18 | 0.095 |
| Cas-Type IE 2 | 28 | 0.004 | | CAS 2 | 4 | 0.021 |
| Cas-Type IE 3 | 37 | 0.005 | | CAS 3 | 5 | 0.026 |
| Cas-Type IE 4 | 6654 | 0.970 | | CAS 4 | 163 | 0.858 |
| Cas-Type IIC 1 | 47 | 0.041 | | Cas-Type IU 1 | 18 | 0.057 |
| Cas-Type IIC 2 | 1 | 0.001 | | Cas-Type IU 2 | 1 | 0.003 |
| Cas-Type IIC 3 | 16 | 0.014 | | Cas-Type IU 3 | 11 | 0.035 |
| Cas-Type IIC 4 | 1088 | 0.944 | | Cas-Type IU 4 | 284 | 0.904 |
| Cas-Type IB 1 | 68 | 0.055 | | Cas-Type VIB1 1 | 1 | 0.009 |
| Cas-Type IB 2 | 8 | 0.006 | | Cas-Type VIB1 2 | 2 | 0.019 |
| Cas-Type IB 3 | 9 | 0.007 | | Cas-Type VIB1 3 | 1 | 0.009 |
| Cas-Type IB 4 | 1162 | 0.932 | | Cas-Type VIB1 4 | 102 | 0.962 |
| Cas-Type IC 1 | 30 | 0.020 | | Cas-Type IIB 1 | 0 | 0.000 |
| Cas-Type IC 2 | 23 | 0.016 | | Cas-Type IIB 2 | 1 | 0.020 |
| Cas-Type IC 3 | 20 | 0.014 | | Cas-Type IIB 3 | 1 | 0.020 |
| Cas-Type IC 4 | 1399 | 0.950 | | Cas-Type IIB 4 | 47 | 0.959 |
| Cas-Type IF 1 | 15 | 0.009 | | Cas-Type VA 1 | 1 | 0.024 |
| Cas-Type IF 2 | 0 | 0.000 | | Cas-Type VA 2 | 0 | 0.000 |
| Cas-Type IF 3 | 7 | 0.004 | | Cas-Type VA 3 | 1 | 0.024 |
| Cas-Type IF 4 | 1625 | 0.987 | | Cas-Type VA 4 | 40 | 0.952 |
| Cas-Type IIIB 1 | 49 | 0.078 | | Cas-Type VIA 1 | 3 | 0.333 |
| Cas-Type IIIB 2 | 0 | 0.000 | | Cas-Type VIA 2 | 0 | 0.000 |
| Cas-Type IIIB 3 | 20 | 0.032 | | Cas-Type VIA 3 | 2 | 0.222 |
| Cas-Type IIIB 4 | 558 | 0.890 | | Cas-Type VIA 4 | 4 | 0.444 |
| Cas-Type IA 1 | 34 | 0.080 | | Cas-Type IIIC 1 | 11 | 0.212 |
| Cas-Type IA 2 | 1 | 0.002 | | Cas-Type IIIC 2 | 0 | 0.000 |
| Cas-Type IA 3 | 11 | 0.026 | | Cas-Type IIIC 3 | 0 | 0.000 |
| Cas-Type IA 4 | 377 | 0.891 | | Cas-Type IIIC 4 | 41 | 0.788 |
| Cas-Type IIA 1 | 4 | 0.005 | | Cas-Type IV 1 | 7 | 0.259 |
| Cas-Type IIA 2 | 0 | 0.000 | | Cas-Type IV 2 | 4 | 0.148 |
| Cas-Type IIA 3 | 14 | 0.016 | | Cas-Type IV 3 | 1 | 0.037 |
| Cas-Type IIA 4 | 855 | 0.979 | | Cas-Type IV 4 | 15 | 0.556 |
| Cas-Type IIIA 1 | 47 | 0.035 | | Cas-Type VB 1 | 0 | 0.000 |
| Cas-Type IIIA 2 | 21 | 0.016 | | Cas-Type VB 2 | 0 | 0.000 |
| Cas-Type IIIA 3 | 23 | 0.017 | | Cas-Type VB 3 | 0 | 0.000 |
| Cas-Type IIIA 4 | 1239 | 0.932 | | Cas-Type VB 4 | 12 | 1.000 |
| Cas-Type ID 1 | 19 | 0.113 | | Cas-Type VIC 1 | 5 | 0.714 |
| Cas-Type ID 2 | 0 | 0.000 | | Cas-Type VIC 2 | 0 | 0.000 |
| Cas-Type ID 3 | 1 | 0.006 | | Cas-Type VIC 3 | 0 | 0.000 |
| Cas-Type ID 4 | 148 | 0.881 | | Cas-Type VIC 4 | 2 | 0.286 |
| Cas-Type IIID 1 | 30 | 0.083 | | Cas-Type VIB2 1 | 0 | 0.000 |
| Cas-Type IIID 2 | 2 | 0.006 | | Cas-Type VIB2 2 | 0 | 0.000 |
| Cas-Type IIID 3 | 9 | 0.025 | | Cas-Type VIB2 3 | 1 | 0.200 |
| Cas-Type IIID 4 | 320 | 0.886 | | Cas-Type VIB2 4 | 4 | 0.800 |

**Table S3. Performance of the machine learning-based classification module in CRISPR-Cas-Docker.** We used K-Nearest Neighbors (K=1) with Hamming distance for the model. The dataset consisted of 16,972 crRNA sequences, with 80% of the data used for training and 20% for testing. The Support column indicates the number of actual instances of the Type in the test set. We pad the shorter sequence with padding characters in order to equalize their lengths when using the Hamming distance measure.

| Hamming nearest neighbor classification | | | | | |
|---|---|---|---|---|---|
| Amount | Type | Precicion | Recall | F1score | Support |
| Major >1,000 | IE | 0.990 | 1.000 | 0.990 | 1373 |
| | IIC | 0.950 | 0.980 | 0.960 | 230 |
| | IB | 0.820 | 0.890 | 0.850 | 249 |
| | IC | 0.940 | 0.960 | 0.950 | 294 |
| | IF | 0.970 | 1.000 | 0.990 | 330 |
| | IIIA | 0.910 | 0.860 | 0.880 | 266 |
| Minor >300 | IIIB | 0.690 | 0.540 | 0.610 | 125 |
| | IA | 0.820 | 0.890 | 0.850 | 85 |
| | IIA | 0.980 | 1.000 | 0.990 | 174 |
| | IIID | 0.680 | 0.580 | 0.630 | 72 |
| Tiny <300 | ID | 0.650 | 0.820 | 0.730 | 34 |
| | VIB1 | 1.000 | 0.900 | 0.950 | 21 |
| | IIB | 0.890 | 0.800 | 0.840 | 10 |
| | VA | 0.830 | 0.620 | 0.710 | 8 |
| | VIA | 1.000 | 0.500 | 0.670 | 2 |
| | IIIC | 0.500 | 0.100 | 0.170 | 10 |
| | IV | 0.000 | 0.000 | 0.000 | 6 |
| | VB | 0.000 | 0.000 | 0.000 | 2 |
| | VIC | 1.000 | 1.000 | 1.000 | 2 |
| | VIB2 | 0.500 | 1.000 | 0.670 | 1 |
| Undefined | CAS | 0.410 | 0.420 | 0.420 | 38 |
| | IU | 0.930 | 0.860 | 0.890 | 63 |
| Macro avg | | 0.750 | 0.720 | 0.720 | 3395 |
| Weighted avg | | 0.920 | 0.930 | 0.920 | 3395 |
| Accuracy | | 0.930 | | | 3395 |
